# Mutation signatures inform the natural host of SARS-CoV-2

**DOI:** 10.1101/2021.07.05.451089

**Authors:** Shanjun Deng, Ke Xing, Xionglei He

## Abstract

The before-outbreak evolutionary history of SARS-CoV-2 is enigmatic because it shares only ∼96% genomic similarity with RaTG13, the closest relative so far found in wild animals (horseshoe bats). Since mutations on single-stranded viral RNA are heavily shaped by host factors, the viral mutation signatures can in turn inform the host. By comparing publically available viral genomes we here inferred the mutations SARS-CoV-2 accumulated before the outbreak and after the split from RaTG13. We found the mutation spectrum of SARS-CoV-2, which measures the relative rates of 12 mutation types, is 99.9% identical to that of RaTG13. It is also similar to that of two other bat coronaviruses but distinct from that evolved in non-bat hosts. The viral mutation spectrum informed the activities of a variety of mutation-associated host factors, which were found almost identical between SARS-CoV-2 and RaTG13, a pattern difficult to create in laboratory. All the findings are robust after replacing RaTG13 with RshSTT182, another coronavirus found in horseshoe bats with ∼93% similarity to SARS-CoV-2. Our analyses suggest SARS-CoV-2 shared almost the same host environment with RaTG13 and RshSTT182 before the outbreak.

## Introduction

Darwin’s evolutionary theory has been challenged ever since it was proposed by the unavailability of some key intermediates between extant species^1^. Importantly, the growing understanding of life in the past one and half century, particularly since the time of molecular biology, provided indisputable intermediate-free supports to Darwin’s theory. When we examine the genomes of current human and, say, chimpanzee, mouse, fish and fly, it’s clear that the delicate principles operating in the non-human species apply to humans as well. There is simply no need to call for a special creator or designer to explain the origin of human beings.

Today we are facing a similar scenario Darwin used to face. The debate on the natural or unnatural origin of SARS-CoV-2, the causative virus of COVID-19, has existed since the beginning of the outbreak^2^ and surged lately^3,4^. One of the main reasons is that RaTG13, the closest relative so far found^5^ (in horseshoe bats *Rhinolophus affinis*), has only ∼96% nucleotide similarities to SARS-CoV-2 (with ∼1,200 nucleotide differences). The situation is distinct from the two previous coronavirus outbreaks happened this century (SARS at 2003 and MERS at 2012); in both cases, a closely related virus with over 99% nucleotide similarities to the causative virus was found in wild animals shortly after the start of each outbreak^6,7^. The missing intermediates between RaTG13 and SARS-CoV-2 prevent a better understanding of the spillover. Fortunately, the signatures left on the available viral genomes would inform the before-outbreak history of SARS-CoV-2.

SARS-CoV-2 belongs to the Betacoronavirus genus, with a single-stranded positive-sense RNA genome of ∼30 thousand nucleotides^8^. There are 12 types of substitution mutations on the viral genome: C>U, C>A, C>G, G>U, G>A, G>C, A>U, A>G, A>C, U>A, U>G, and U>C. The genome-wide mutation spectrum, which measures the relative rates of the 12 mutation types, comprises a set of summary statistics with little functional relevance. More importantly, the viral mutation spectrum is expected to be heavily shaped by host factors^9^. For example, the large number of RNA-binding proteins in mammalian cells would necessarily interact with the single-stranded RNA genome^10^, which is critical for preventing the hydrolytic deamination of cytosines (leading to C>U) and the reactive oxygen species (ROS) induced oxidation of guanines (leading to G>U)^11^. Also, the two key RNA editing protein families, ADAR^12^ (adenosine deaminase acting on RNA) and APOBEC^13^ (apolipoprotein B mRNA editing enzyme catalytic polypeptide-like), would cause A>G and C>U mutations, respectively. In addition, when the host immunity failed to prevent high virion production, the cellular supply of dATP, dUTP, dCTP, and dGTP would modulate the viral mutations during genome replication^14^. The activities of the host factors often vary substantially among different species or even among different tissues of the same species^15^, and their interplay would be even more complex. Hence, the viral mutation spectrum as a 12-dimension signature vector would be a powerful tool for tracking the hosts.

## Results

### Evolution of mutation spectrum in the SARS-CoV-2 lineage

We included SARS-CoV-2 and six related viruses in the analysis (Fig. 1a). The six related viruses were chosen because they are evolutionarily close enough for reliable mutation inferences while distant enough for observing plenty of mutations. At least three different hosts, bat, pangolin and human, are involved, highlighting a complex host history of this viral lineage^16,17^. Two separate phylogenetic trees were constructed to avoid the phylogeny confusions caused by recombination (Fig. S1), which results in different genealogical histories at different genomic regions in the ancestor of Bat-Cov-ZXC21 and Bat-Cov-ZC45 (both found in horseshoe bats *Rhinolophus sinicus*^18^). The branch X, which represents the before-outbreak history of SARS-Cov-2, and the B1, which represents the history of RaTG13 after it split from SARS-Cov-2, are present in both phylogenetic trees. Using conventional molecular evolutionary methods^11^, we compared the viral genomes to infer the substitution mutations occurred on the evolutionary branches as marked in Fig. 1a (Methods). We considered only the third codon positions such that the obtained mutation spectra are less shaped by selection^19^ (Fig. 1b and Table S1). Because the mutations on different evolutionary branches occurred independently, the derived mutation spectra of the branches are independent. To quantify the similarity between two mutation spectra we computed an identity score (i-score), which is the proportion of the total rate variation explained by the x=y dimension in a two-dimensional plot of the two spectra as in Fig. 1c (Methods). An i-score equal to 100% means the two mutation spectra are 100% identical.

**Fig. 1.**
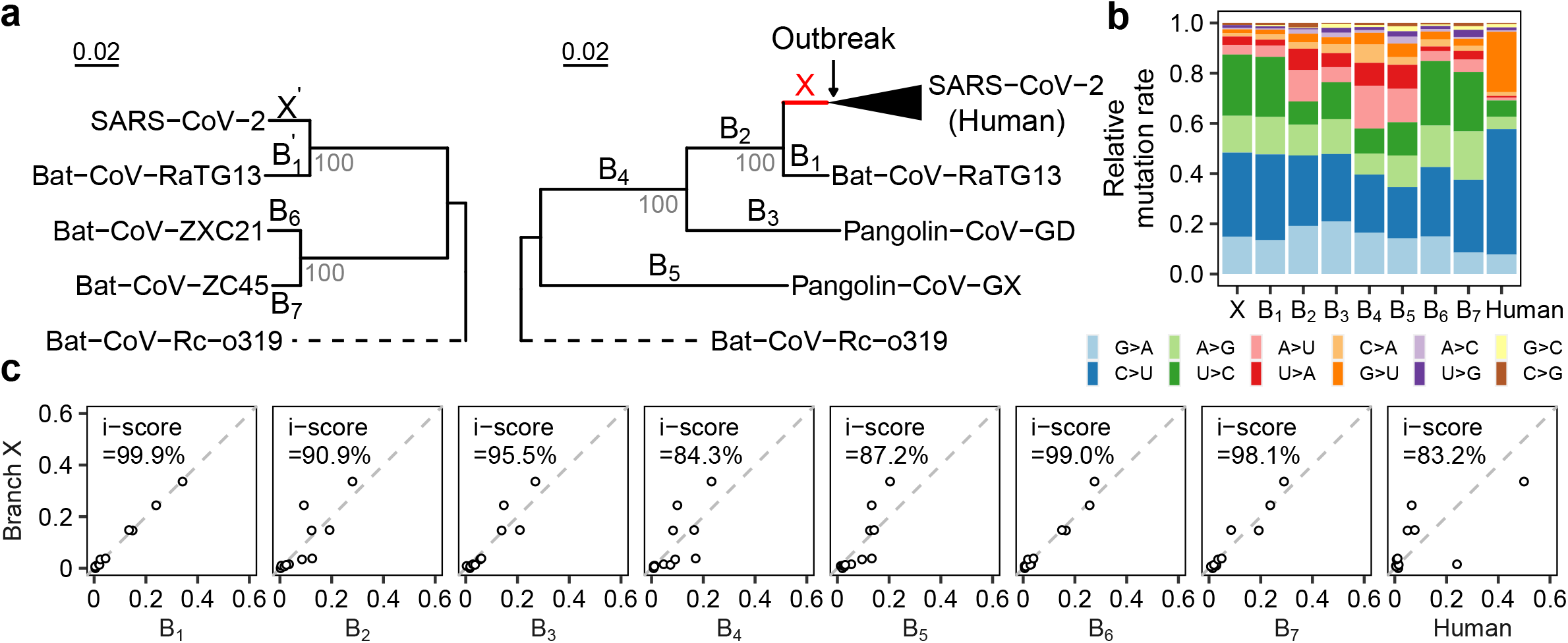
Evolution of mutation spectrum in the SARS-CoV-2 lineage. **a**. The phylogenetic relationships of the seven coronaviruses included in the analysis. Two separate phylogenetic trees are considered to resolve the confusions caused by recombination, which results in different genealogical histories at different genomic regions in the ancestral branch of Bat-CoV-ZXC21 and Bat-CoV-ZC45. Nine major evolutionary branches examined in this study, X, B1-B7, and the Human branch, are shown. The branch X and B1 are also present (as X’ and B1’) in the tree with B6 and B7 to help infer the ancestor of B6 and B7. The Bat-CoV-Rc-o319 is used as outgroup in both trees. **b**. The relative mutation rate of the 12 mutation types on each of the nine evolutionary branches. **c**. The similarity of mutation spectrum between branch X and each of the other eight branches. The similarity of two branches is measured by identity score (i-score), which is the proportion of total rate variation explained by the x=y dimension in the plot of the two spectra.

The mutation spectra calculated separately in the two phylogenetic trees are nearly identical for the same branches (i-score = 99.9% for X versus X’ and 99.4% for B1 versus B1’; Fig. S2), suggesting the results of the two trees comparable. There are three notable features regarding the obtained spectra (Fig. 1b-c). First, the branch X is nearly identical to B1, with an i-score = 99.9%. Second, the branch X is distinct from the after-outbreak branch of SARS-CoV-2 (i.e, the Human branch), with an i-score = 83.9%. The obtained spectrum of the Human branch is consistent with a previous study^9^. Compared to branch X, the Human branch has a lot more G>U and C>U mutations, suggesting much stronger mutational pressures imposed by ROS and APOBEC family, respectively, to the SARS-CoV-2 genome in infected human cells. Meanwhile, the rates of A>G/U>C mutations reduce substantially, suggesting weaker activity of the ADAR family. Third, the branch X is in general highly similar to the branches with bats as the putative hosts (B1, B6 and B7) while less similar to the branches with non-bat hosts involved. These results, in particular, the 99.9% identify of X and B1, suggest SARS-CoV-2 not be artificially synthesized for gain-of-function research, because mutation spectrum is of little functional relevance and a synthesized genome is unlikely to show such a similar mutation spectrum to a naturally evolved viral genome (RaTG13). Notably, making comparably similar mutation spectra is doable by nature for close sister lineages like B6 and B7 (Fig. S2)

### Host signatures inferred from viral mutations

The viral mutations are caused by both replication errors and replication-independent lesions or editing. The former is mostly associated with the viral self-encoded replication-transcription complex (RTC) and the latter would be mostly explained by host factors^20^ (Fig. 2a). The coronavirus positive-sense RNA genome is replicated first by forming a negative-sense RNA intermediate, which then serves as template for both transcription and replication^8^. The same replication errors occurred in producing negative-sense strand and in producing positive-sense strand would result in different mutation types. For example, the two steps for replicating a nucleotide C (C-to-G followed by G-to-C) are the same, but in an opposite order, as the two steps of replicating a G (G-to-C followed by C-to-G). Then, the same replication error of, say, C-to-A, in the C-to-G step would cause a C>U mutation in the replication of C but a G>A mutation in the replication of G (Fig. 2a). Other types of replication errors have the same feature. As a result, the 12 mutation types would form six complementary pairs: C>A/G>U, C>U/G>A, C>G/G>C, A>U/U>A, A>C/U>G, and A>G/U>C; in each pair the two complementary mutation types would have the same rate if all mutations were due to replication errors. Hence, the different mutation rate observed in each complementary pair would be ascribed to replication-independent factors, which are associated in a large part with host. For example, the preferential binding of the host APOBEC family to the single-stranded positive-sense RNA would lead to more C>U mutations than G>A mutations^21^. The host ADAR family would preferentially edit the negative-sense strand that are often in a double-stranded form, resulting in more U>C mutations than A>G mutations^22^. In addition, the damage effects of ROS primarily on single-stranded RNA would cause a higher rate of G>U mutations over C>A mutations^23^. The direction and magnitude of the rate difference in each complementary pair then constitute a signature of host factors, which informs the identity of hosts.

**Fig. 2.**
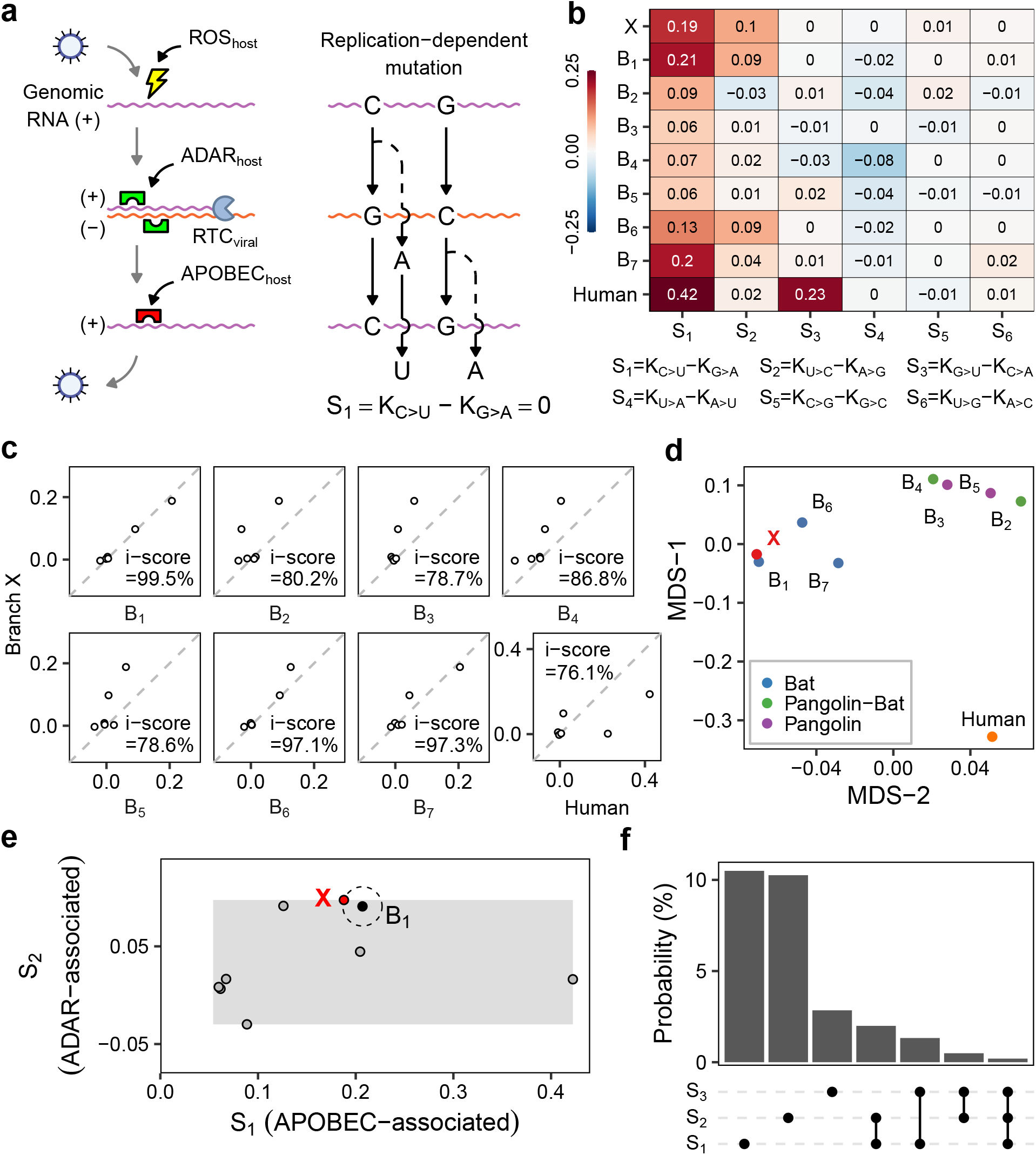
Host signatures inferred from viral mutation spectrum. **a**. A diagram showing the major sources of viral mutations, which include the replication errors (by the viral replication-transcription complex RTC) and the lesions caused by host factors. Because replication processes are the same, despite in the opposite order, for nucleotides G and C (or A and T), replication errors would result in equal rates of complementary mutations such as C>U and G>A. However, host factors would distort the equal-rate pattern of complementary mutation pairs. The positive-sense RNA is often in a single-stranded form, sensitive to ROS and the APOBEC family, while the negative-sense RNA tends to be in a double-stranded form, thus more affected by the ADAR family. **b**. The rate difference of each complementary mutation pair serves as a signature of host factors. There are thus six host signatures, each corresponding to a complementary mutation pair, inferred from the viral mutation spectrum. Among the three major host signatures, S1 is likely associated with the APOBEC family, S2 the ADAR family, and S3 the ROS. **c**. The similarity of host signatures between branch X and each of the other eight branches. Branch X is highly similar to B1, B6 and B7, the three branches of bat coronavirus. **d**. A multidimensional scaling (MDS) plot of the host signatures reveals nearly the same positions of branch X and B1. **e**. Estimation of the likelihood that an arbitrary laboratory condition happens to match the host signatures of B1 (the branch of RaTG13). The grey rectangle area is defined by the empirical ranges of S1 (APOBEC-associated) and S2 (ADAR-associated) that are based on the data of panel b. The probability of approaching B1 as closely as X is the area of the circle divided by the whole rectangle area, which is ∼2.0%. The positions of the other seven branches are also shown in the rectangle area. **f**. The probability that an arbitrary condition approaches B1 as closely as X is given, by considering the different combinations of S1, S2, and S3, respectively.

To obtain the host signatures we calculated the rate difference in each complementary pair. The six host signatures (S1-S6), each corresponding to a complementary pair, are indeed informative (Fig. 2b). For example, S1, the rate of C>U minus the rate of G>A, ranges from 0.06 to 0.42 among the different evolutionary branches. This may represent the different activities of the APOBEC family in different hosts. S2, the rate of U>C minus the rate of A>G, ranges from −0.03 to 0.1. This is likely associated with the relative activity of the ADAR family. S3, the rate of G>U minus the rate of C>A, ranges from −0.03 to 0.23 and appeared unusually strong in the Human branch. This could be related to ROS that may preferentially target the single-stranded positive-sense RNA and have a strong induction in the infected human cells. Notably, the mentioned genes/pathways are just putatively associated with the observed host signatures. We found branch X has nearly identical host signatures to B1, with an i-score = 99.5%, despite substantial deviations from the human or pangolin associated branches (Fig. 2c). A multidimensional scaling plot shows that X is almost perfectly overlapping with B1, close to B6 and B7, and distant from the other branches (Fig. 2d). These results suggest that SARS-CoV-2 shared almost the same host environment with RaTG13 before the outbreak.

To gauge the probability that an arbitrary cell culture condition in laboratory matches the natural host environment of RaTG13, we estimated the size of the space formed by the host signatures, each of which has an empirical range according to the nine branches presented in Fig. 2b. We considered S1, S2 and S3 because their empirical ranges are the largest and their associated genes/pathways (APOBEC, ADAR and ROS) appear independent. As shown in Fig. 2e, the probability of approaching, as closely as SARS-CoV-2, the host environment of RaTG13 is ∼2.0%, if S1 and S2 are considered. The number would be 0.02% if S3 is also considered (Fig. 2f). The estimations are conservative because the other three signatures (S4-S6) were not considered and also the real ranges of the signatures would be larger than the empirical ranges based on the nine evolutionary branches. We cautioned that the calculations assumed the associated gene/pathway activities are uniformly distributed within the empirical ranges. Nevertheless, the results are helpful for thinking of the likelihood that an arbitrary cell culture condition set in laboratory happens to duplicate a defined natural host environment.

### Robust signals after replacing RaTG13 with RshSTT182

Because there are concerns on the quality of the assembled genome of RaTG13^24^, we reproduced the above analyses after replacing RaTG13 with another bat coronavirus RshSTT182. RshSTT182 was isolated from Shamel’s horseshoe bats (*Rhinolophus shameli*), being the first close relative of SARS-CoV-2 found in Southeast Asia (Cambodia) and with 92.6% genomic identity to SARS-CoV-2^25^. The whole-genome phylogeny of the involved viruses is (((((SARS-CoV-2, RaTG13), RshSTT182), Pangolin-CoV-GD), Pangolin-CoV-GX), Rc-o319). Hence, replacing RaTG13 with RshSTT182 would affect mainly the branches X, B1, and B2 in our analyses. Using the same procedure we obtained the mutation spectra and derived the host signatures for each of the evolutionary branches. The findings remain qualitatively the same (Fig. S3-S4 and Table S2). In brief, the mutation spectrum of SARS-CoV-2 is 99.3% identical to that of RshSTT182 (99.9% in the case of RaTG13). The slight reduction of the similarity may reflect the fact that the host of RaTG13 is *Rhinolophus affinis* but the host of RshSTT181 is another horseshoe bat species *Rhinolophus shameli*. Taken together, our analyses suggest the host environment of SARS-CoV-2 before the outbreak be fully compatible with horseshoe bats.

## Discussion

It should be emphasized that this study is to address the evolution of the SARS-CoV-2 genome but nothing else. Using mutational signatures inferred from the available viral genomes we probed the evolutionary time window (branch X) SARS-CoV-2 spent before the outbreak and after the split from bat coronavirus RaTG13. The missing intermediates within this time window that presumably spans a few tens of years^26^ prevents a better understanding of the spillover. Our analyses based on public data provide compelling evidence that during this time window SARS-CoV-2 evolved in a host environment highly similar, if not identical, to RaTG13. The host environment is also similar to that of the three bat coronaviruses RshSTT182, ZXC21 and ZC45, and difficult to duplicate by an arbitrary cell culture condition set in laboratory. One may argue that, while the branch X as a whole is compatible with natural laws, it may not be at a few key sites. Such an argument presumes that there are intermediates with over 99% similarity to SARS-CoV-2 to be found in nature. Notably, claiming such natural intermediates would leave little room for speculations, as in the cases of SARS^6^ and MERS^7^. The mission of the scientific community is then to find them in nature to better understand the spillover.

## Methods

### Genomic Data

The SARS-CoV-2 related bat and pangolin coronavirus genomic sequences were obtained from NCBI GenBank (https://www.ncbi.nlm.nih.gov/genbank). For genomes without accurate annotations of ORFs, we re-annotated these genomes with CDSs annotated in SARS-CoV-2 by Exonerate2 (–model protein2genome: bestfit –score 5 -g y)^27^. The complete genomic sequences and metadata of SARS-CoV-2 were retrieved from Global Initiative on Sharing All Influenza Data (GISAID; https://www.gisaid.org/; accessed on 19 March 2021)^28^. Gap-containing genomes in examined regions were removed, and only genomes from Dec. 2019 to Dec. 2020 were chosen for analysis. All available genomes submitted to GISAID from Dec. 2019 to Feb. 2020 were included, and, among the too many submitted genomes from Mar. to Dec. 2020, 2,000 genomes were randomly selected for each month. Finally, a total of 214,32 SARS-CoV-2 genomes were included. Following GISAID we used SARS-CoV-2 WIV04 (EPI_ISL_402124) as the reference genome. The detailed information of SARS-CoV-2 and the related coronaviruses included in this analysis is summarized in Supplementary Dataset I.

### Phylogenetic analysis and mutation spectra calculation

The codon alignments of ORFs were performed based on amino acid sequences translated by TranslatorX^29^ and MAFFT v7.471^30^, and further concatenated by AMAS^31^ and refined with visual check. Only ORFs with consistent annotations in the examined viruses were included. Maximum likelihood phylogenetic analysis based on the whole coding regions was conducted by using IQ-TREE v2.0.3^32^ with GTR+FO+R10 substitution model and 1,000 bootstrap replicates. The ancestral sequences of the internal nodes were inferred in IQ-TREE with an *-asr* parameter, and mutations on each branch were derived by comparing the ancestral sequence to the descendant sequence. To avoid the confounding effects of potential recombination and convergent evolution, the region covering the receptor binding domain and the furin-like cleavage site (319^th^-770^th^ codons) of the spike protein was removed from the analysis. Only the third codon positions were considered in calculation of the mutation spectra. The aligned sequences can be found in Supplementary Dataset II-V.

To obtain the after-outbreak mutations of SARS-CoV-2, 59 separate main clades each containing more than 100 sequences and supported by a bootstrap value >90 were selected from the phylogenic tree. Mutations were inferred by comparing each individual sequences to the corresponding common ancestral sequences of each clade, respectively. To avoid redundancy, recurrent mutations within a clade were counted once. Then, the 59 clade-specific ancestral sequences were compared to the earliest common ancestral sequence of SARS-CoV-2. Mutations obtained from the two steps were pooled to derive the mutation spectrum of the Human branch.

For a specific mutation type, say C>A, the rate was calculated as the number of C>A mutations divided by the total number of C nucleotides in the ancestral sequence of the given branch (third codon positions). The mutation rates of the 12 mutation types were then each divided by their sum to obtain the relative mutation rates (i.e., mutation spectrum). The i-score of two mutation spectra is the proportion of variance explained by the x=y dimension in a two-dimensional plot of the two spectra. Specifically, let A = [S_1_, S_2_]^*T*^, where S_1_ and S_2_ are the two mutation spectra under examination, and B = [D_1_, D_2_]^*T*^, where D_1_ is the projection of A onto the x=y dimension and D_2_ onto the x= -y dimension. Then, the i-score = cov(D_1_) / (cov(S_1_)+ cov(S_2_)).

To verify the whole-genome-based evolutionary branches at different genomic regions a sliding window analysis through the viral genomes was conducted. Specifically, each window covers 500 codons (or 1500 nucleotides, ∼5% of the viral genome) and the step size is a half window. For each window we constructed the phylogeny of the viruses using synonymous sites, and then checked if the whole-genome-based branches exist in the window. Neighbor-Joining phylogeny was obtained in MEGA X^33^, which allows such analysis on synonymous sties, with 1,000 bootstrap replicates.

